# Co-circulating TBEV lineages and evidence for vertical transmission in wild bank voles from Poland

**DOI:** 10.64898/2026.07.27.741075

**Authors:** Martyna Krupińska, Teemu Smura, Lukasz Rąbalski, Katarzyna Tolkacz, Beata Biernat, Dorota Dwużnik, Karolina Baranowicz, Sanna Maki, Martyna Krejmer-Rabalska, Klaudia Baranska, Ravi Kant, Tarja Sironen, Olli Vapalahti, Jerzy M. Behnke, Anna Bajer, Maciej Grzybek

## Abstract

Tick-borne encephalitis virus (TBEV) is a zoonotic flavivirus maintained in complex enzootic cycles involving ticks and vertebrate hosts. While vector-mediated transmission has been extensively studied, host-associated mechanisms, including vertical transmission and prolonged viral carriage in natural reservoir populations, remain poorly understood. Here, we investigated TBEV circulation, genetic diversity, and evidence for vertical transmission in bank voles (*Clethrionomys glareolus*) from a highly endemic region of North-eastern Poland. A total of 258 wild rodents were screened using molecular and serological approaches. TBEV RNA was detected in 14.3% of individuals, whereas seroprevalence was substantially lower (6.6%), revealing limited concordance between viral presence and humoral response under natural conditions. Near-complete genome sequences were obtained from 10 TBEV-positive individuals, representing the first near-complete TBEV genomes generated from wild rodents in Poland. All isolates belonged to the European subtype (TBEV-Eu), and phylogenetic analysis revealed two genetically distinct viral clades co-circulating within a narrow spatiotemporal window. Importantly, TBEV RNA and/or envelope protein were detected in embryos from naturally infected females, providing evidence consistent with vertical transmission in a wild reservoir host. Together, these findings suggest that vertical transmission and complex host infection dynamics may be underappreciated components of TBEV maintenance in natural reservoir populations. Our results highlight the need to integrate vertebrate host dynamics into models of TBEV ecology and support expanded wildlife-based surveillance to better understand and predict zoonotic risk.

## Introduction

Tick-borne encephalitis virus (TBEV) is a widespread Eurasian orthoflavivirus capable of causing infections of the central nervous system in humans and other vertebrates[1]. Clinical manifestations range from asymptomatic infections to severe encephalitis, with marked differences in severity among the three major subtypes: European (TBEV-Eu), Siberian (TBEV-Sib), and Far Eastern (TBEV-FE) [2–5].

In natural ecosystems, TBEV circulates in complex transmission networks involving ixodid ticks (principally *Ixodes ricinus* and *I. persulcatus*) and vertebrate reservoir hosts [1]. Rodents play a central role in supporting both viremic and non-viremic transmission and harboring latent infections (Figure 1) Co-feeding transmission enables efficient virus transfer even without detectable viremia, allowing transmission despite pre-existing host antibodies [6, 7]. However, key aspects of infection dynamics, including persistence, immune responses, and non-vectorial transmission, remain poorly understood.

**Figure 1.**
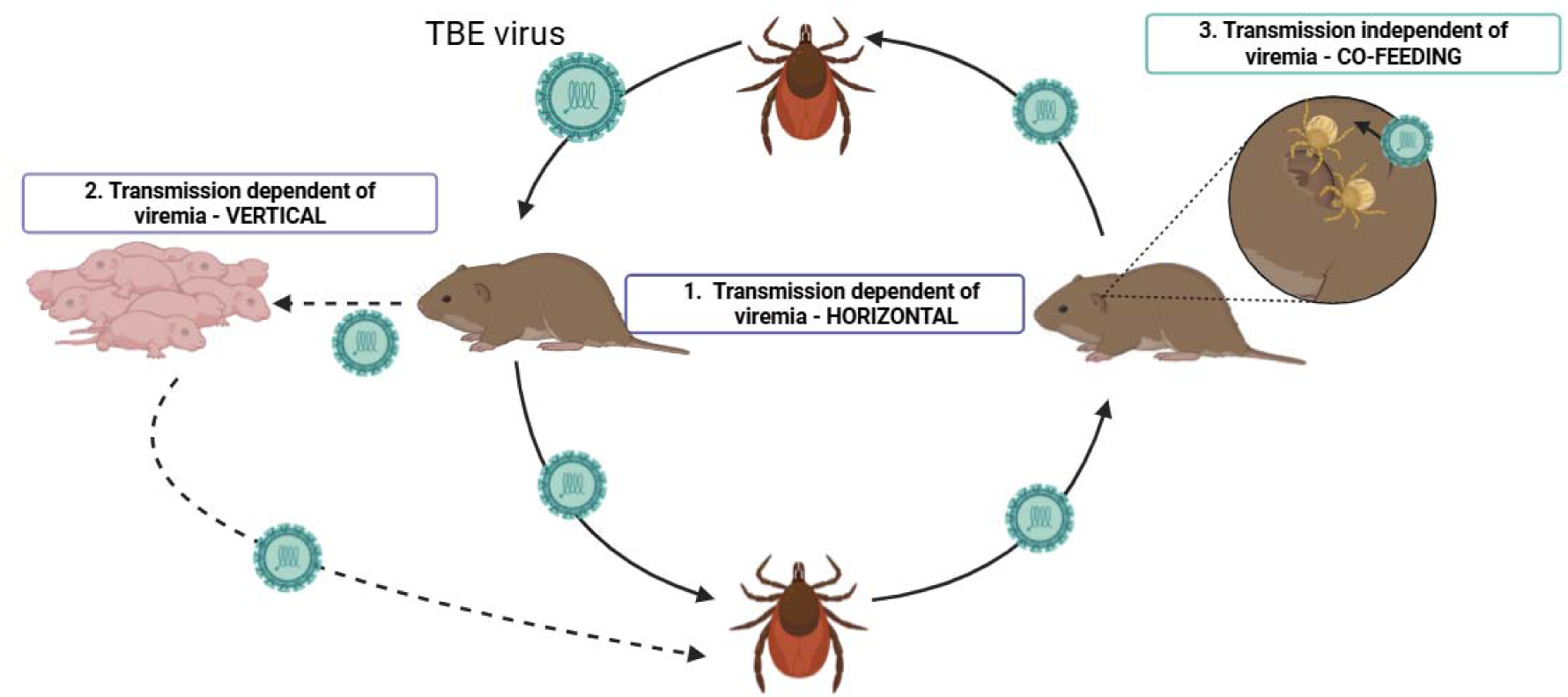
Transmission pathways of TBEV in rodent hosts. TBEV circulation in rodents occurs via both viremic and non-viremic transmission routes. (1) Following infection through a tick bite, rodents may develop viremia, enabling transmission to naïve ticks during blood feeding (horizontal transmission). (2) Vertical transmission may occur potentially from infected females to offspring via transplacental transfer and/or through lactation. (3) Co-feeding transmission occurs when infected and uninfected ticks feed simultaneously in proximity on the same host, enabling localized virus transfer through the skin without requiring systemic viremia and even in the presence of circulating host antibodies. Created with BioRender.com.

Traditional tick surveillance has limited sensitivity due to low infection prevalence and spatial heterogeneity in virus distribution [8]. Rodents may provide a more integrative measure of local virus circulation [6, 7, 9–11].

Vertical transmission among mammalian hosts represents a particularly understudied mechanism of TBEV maintenance. While experimental studies have demonstrated transplacental and sexual transmission of TBEV-FE in a non-adapted host - laboratory mice[12], evidence from a natural reservoir system is restricted to red-backed voles (*Clethrionomys rutilus*) [13]. In the latter study, the route of mother-to-offspring transmission could not be determined, as both transplacental and lactational transmission were possible. Whether such transmission occurs for TBEV-Eu in wild rodent populations remains unknown.

Given the rising TBE incidence and geographic expansion in Europe, improved understanding of transmission mechanisms is essential [14–23]. Poland represents a potential zone of subtype coexistence, yet molecular data on circulating strains remain limited [24, 25].

Here, we investigated TBEV circulation in bank voles (*Clethrionomys glareolus*) from a highly endemic region of North-eastern Poland. Specifically, we aimed to (i) determine molecular prevalence and seroprevalence, (ii) characterize the genetic diversity and phylogenetic relationships of circulating viral strains, and (iii) assess evidence for vertical transmission in a natural reservoir host. We hypothesized that host-associated transmission contributes to the maintenance of TBEV in natural reservoir populations and tested this using integrated molecular, serological, and phylogenomic analyses of wild bank voles.

## Materials and methods

A total of 258 bank voles were sampled from three ecologically comparable sites in North-eastern Poland. Blood samples were collected via cardiac puncture and screened for TBEV-specific antibodies using an indirect immunofluorescence assay. Brain tissues were obtained during necropsy, and total RNA was extracted for molecular detection of TBEV using RT-nested PCR. Embryos from TBEV-positive pregnant females were additionally examined for the presence of viral RNA and TBEV envelope antigen. RNA extracted from PCR-positive brain samples was subjected to sequencing, and the resulting viral genomes were analyzed using phylogenetic approaches.

### Study sites

The study was conducted at three woodland sites located in the Mazury Lake District in North-eastern Poland (Figure 2): Urwitałt (53°48.2′N, 21°39.7′E), Tałty (53°53.6′N, 21°33.0′E), and Pilchy (53°42.2′N, 21°48.5′E). These sites are separated by natural barriers, including lakes, which may limit direct host movement over short ecological timescales. However, the bank vole population in this region is considered largely panmictic, with previous genetic studies indicating ongoing gene flow among sites [26]. Detailed descriptions of habitat characteristics and study design have been reported previously [27].

**Figure 2.**
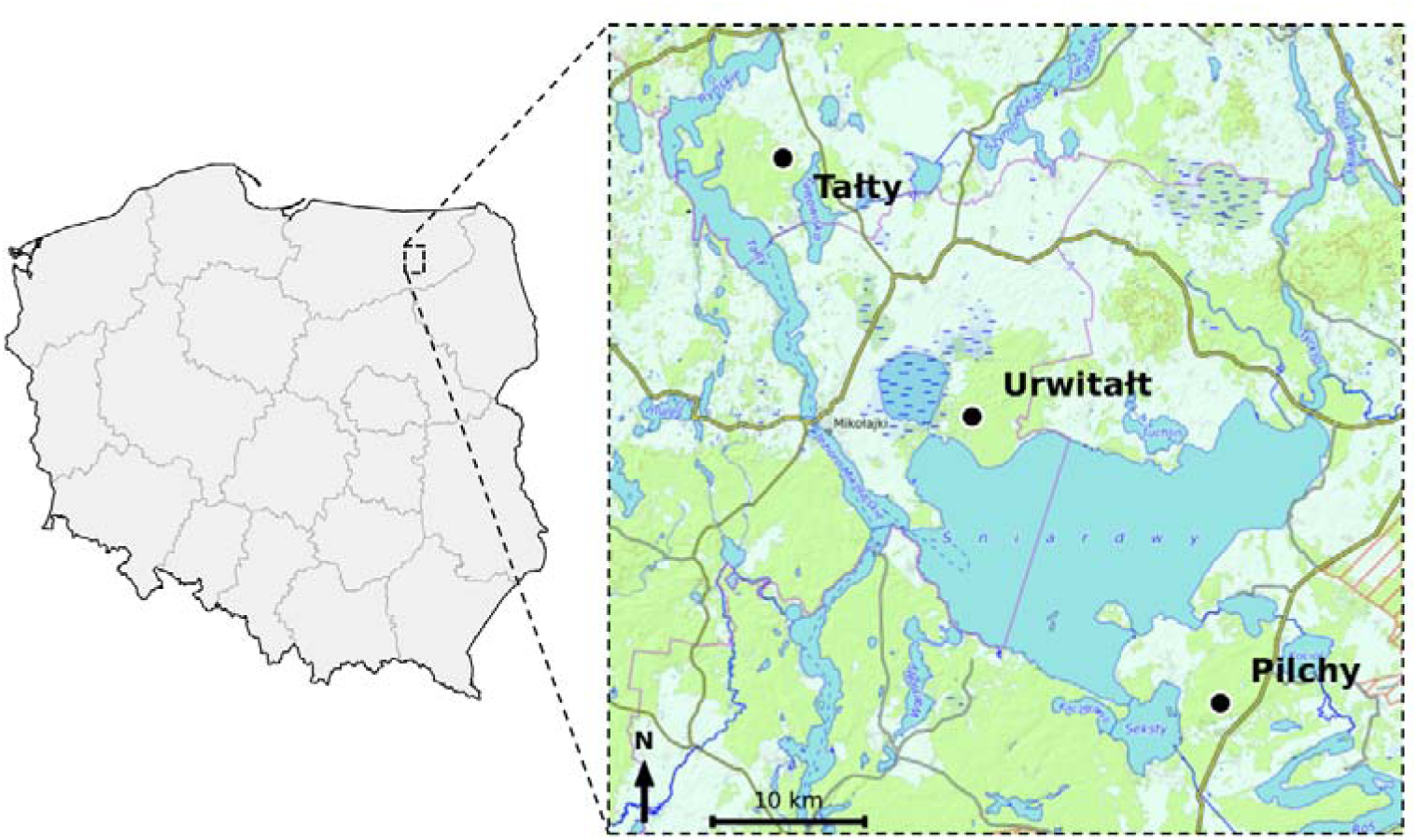
Map with study sites marked, located in northeastern Poland. Rodents were trapped in woodlands near Tałty, Urwitałt and Pilchy. Map data ©OpenTopoMap contributors.

### Collection of rodents

Rodents were sampled between mid-August and mid-September 2018. Trapping was conducted for 3-4 consecutive days at each site. Detailed trapping procedures, as well as animal handling and processing protocols, have been described previously[27–29].

Individuals were assigned to three age classes based on principal component analysis of morphological traits, including body mass and dried eye lens weight: class 1 (immature juveniles), class 2 (young adults), and class 3 (breeding adults) [27, 30]. Animals were euthanized by anesthetic overdose, and blood samples were collected via cardiac puncture using sterile syringes. Samples were allowed to clot at room temperature and centrifuged at 5,000 rpm for 10 min. Serum was separated and stored at -20 °C during fieldwork and subsequently at -80 °C until analysis.

Brain tissues were collected during necropsy under sterile conditions. Skulls were bisected, and brain samples were divided into two portions and preserved in RNA stabilization solution (StayRNA, A&A Biotechnology, Gdansk, Poland). After 24h at 4°C, samples were stored at -20 °C in the field and transferred to -80 °C for long-term storage.

Embryos from pregnant females were collected during necropsy, immediately frozen at -20°C, and subsequently processed within 2h. Embryos were counted, weighed, and measured, then bisected longitudinally using sterile instruments and stored at - 80 °C for downstream molecular and antigen detection analyses.

### RNA extraction and detection of viral RNA

Total RNA was extracted from brain tissue or half of the embryo using the Total RNA Mini Kit (A&A Biotechnology, Gdansk, Poland) according to the manufacturer’s instructions. RNA was eluted in PCR-grade water and stored at -80 °C until further analysis. Reverse transcription was performed using the QuantiTect Reverse Transcription Kit (Qiagen, Hilden, Germany), including a genomic DNA elimination step (42 °C for 2 min), followed by cDNA synthesis at 42 °C for 15 min. RNA quality and concentration were assessed in a subset of samples using a NanoDrop spectrophotometer (Thermo Fisher Scientific, Waltham, MA, USA).

Detection of TBEV RNA was performed using a nested RT-PCR protocol targeting the 5′ non-coding region and the 5′ terminus of the capsid (C) protein gene, as previously described by Ramelow et al. (1993) [31]. Outer primers (TBE1, TBE2) and inner primers (TBE3, TBE4) were used. PCR reactions were carried out using PCR Mix Plus Green (A&A Biotechnology, Gdansk, Poland) with 0.5 μM of each primer. In the first round, 2 μl of cDNA was used as template, followed by a second round using 2 μl of the initial PCR product. Cycling conditions for both rounds consisted of initial denaturation at 94 °C for 3 min, followed by 40 cycles of 94 °C for 30 s, 37 °C for 40 s, and 72 °C for 30 s, with a final extension at 72 °C for 10 min. Each run included a positive control (RNA extracted from the TBEV-Eu reference strain Kumlinge A52) and a negative control (PCR-grade water).

### Indirect immunofluorescence assay for detection of anti-TBEV antibodies

Serum samples from bank voles were analyzed for TBEV-specific antibodies using a validated indirect immunofluorescence assay (IFA), as previously described [32]. Samples were diluted 1:10 in PBS. TBEV-infected Vero E6 cells were detached with trypsin, mixed with uninfected Vero E6 cells (in a ratio of 1:3), washed with PBS, and air-dried on slide spots. As a background control, uninfected Vero E6 cells were used. After spotting and air-drying, the slides were fixed with acetone and stored dry at −70°C until use. Seropositive human serum was included as a positive control and was detected using a FITC-conjugated anti-human IgG secondary antibody, whereas bank vole sera were detected using a FITC-conjugated rabbit anti-mouse IgG secondary antibody (LGC SeraCare, Milford, MA, USA). Slides were examined under a fluorescence microscope, and images were captured using a ZOE™ fluorescent cell imager (Bio-Rad, Hercules, CA, USA).

### SDS-PAGE Western-blot

Protein extracts were prepared from halves of embryo homogenates (4-20 μl) together with positive and negative controls. Samples were denatured at 100 °C for 15 min in Laemmli buffer and separated on 8% SDS-PAGE precast gels (Bolt 8% Bis-Tris Plus, Thermo Fisher Scientific, Waltham, MA, USA). Electrophoresis was performed in 1× Bolt MES SDS running buffer at 120 V for 1.5 h.

Proteins were transferred onto PVDF membranes using wet transfer overnight. Membranes were blocked in 5% (w/v) non-fat milk in Tris-buffered saline with 0.1% Tween 20 (TBS-T), followed by washing (4 × 5 min) in TBS-T. Membranes were incubated with primary antibodies against TBEV envelope (E) protein (kindly provided by Prof. Matthias Niedrig), followed by goat anti-mouse horseradish peroxidase (HRP)-conjugated secondary antibodies (Jackson ImmunoResearch Laboratories, West Grove, PA, USA), each applied at a 1:2000 dilution for 1 h at room temperature.

Protein bands were visualized using SuperSignal™ West Pico Plus chemiluminescent substrate (Thermo Fisher Scientific, Waltham, MA, USA) and imaged with a ChemiDoc™ Alliance Q9 system (UVITEC). As a negative control, lysate from non-transfected FreeStyle™ 293 cells was used, whereas the positive control consisted of lysate from FreeStyle™ 293 cells expressing TBEV E protein (pcDNA3.1-TBEV-E construct).

### Next-generation sequencing

cDNA was synthesized from RT-PCR–positive RNA samples using the LunaScript RT SuperMix Kit (New England Biolabs, Ipswich, MA, USA) with random hexamers and oligo(dT), followed by tiled-amplicon PCR using the Q5 PCR Kit (New England Biolabs) as previously described by Smura et al. [33]. Sequencing libraries were prepared using the NEBNext Ultra II DNA Library Prep Kit for Illumina (New England Biolabs) and purified using SPRIselect beads (Beckman Coulter Life Sciences, Indianapolis, IN, USA). Libraries were quantified using a Qubit fluorometer (Thermo Fisher Scientific, Waltham, MA, USA) and sequenced on an Illumina NovaSeq 6000 platform using a NovaSeq 6000 SP Reagent Kit v1.5 (500 cycles).

Raw sequence reads were trimmed, quality-filtered, and assembled using fastp [34]and BWA-MEM [35] implemented in the HAVoC pipeline [36]. Newly generated sequences were deposited in GenBank under accession numbers PZ361876–PZ361885.

### Sequence alignment and phylogenetic analysis

Obtained TBEV sequences were compared with sequences retrieved from the NCBI database using BLAST to assess sequence similarity. Sequences representing the Siberian and Far Eastern TBEV subtypes were additionally included in the analysis. Redundant sequences originating from the same country and year and clustering within the same phylogenetic clade were excluded from further analysis. Multiple sequence alignment was performed using MAFFT v7.408 with the global pairwise alignment strategy and a maximum of 1000 iterative refinement rounds to ensure high alignment accuracy [37]. Maximum-likelihood phylogenetic analysis was performed using IQ-TREE v2.0.7[38]. The best-fit nucleotide substitution model was selected using ModelFinder according to the Bayesian Information Criterion and identified as GTR+F+I+G4 [39]. Tree reconstruction was performed with 1000 ultrafast bootstrap replicates[40]. Omsk hemorrhagic fever virus was used as an outgroup.

### Statistical analysis

The statistical approach has been documented comprehensively in our earlier publications [9, 27, 29, 41, 42]

Prevalence (proportion of TBEV RNA-positive individuals) and seroprevalence (proportion of anti-TBEV IgG-positive individuals) were calculated with 95% confidence intervals (CI) using the exact binomial method [43]. Associations between infection status (RNA positive/negative) or serological status (IgG positive/negative) and host characteristics were analysed using maximum-likelihood log-linear contingency table analysis implemented in IBM SPSS Statistics version 21. Initially, full factorial models were fitted, incorporating as factors SEX (2 levels, males and females), AGE (3 levels), and SITE (3 levels, Urwitałt, Tałty, Pilchy). The presence or absence of TBEV RNA (PREVALENCE) and antibodies (SEROPREVALENCE) were considered as binary factors. The importance of each term in interactions involving PREVALENCE/ SEROPREVALENCE in the final model was assessed by the probability that its exclusion would alter the model significantly and these values are given in the text.

### Ethical approval

All procedures involving animals were conducted in accordance with the Guidelines for the Care and Use of Laboratory Animals of the Polish National Ethics Committee for Animal Experimentation and the Polish national law for field studies involving the trapping and culling of unprotected vertebrates for scientific purposes (Resolution No. 12/2022 of the Polish National Ethics Committee on Animal Experimentation, 11 March 2022). Appropriate permits were obtained for field trapping and subsequent laboratory analyses. The study protocol was reviewed and approved by the First Warsaw Local Ethics Committee for Animal Experimentation (approval no. 76/2015). All rodents were culled by Prof. Anna Bajer (authorized to implement experimental procedures and the culling of animals for scientific objectives by the Polish Laboratory Animal Science Association, (License number, 13/2015). All efforts were made to minimize animal suffering, and animals were handled by trained personnel in accordance with established ethical standards. The study is reported in accordance with the ARRIVE guidelines 2.0 [44].

## Results

### Prevalence and seroprevalence of TBEV

TBEV infection was assessed in 258 bank voles using molecular and serological approaches. Viral RNA was detected in brain samples of 37 individuals, corresponding to an overall prevalence of 14.3% (95% CI: 10.6-19.1). The prevalences of PCR positivity for TBEV in all the data subsets and overall are shown in Table 1. Prevalence was very similar in voles from each of the three sites, ranging from 11.8% in Pilchy to 16.1 % in Tałty (χ*^2^_2_*=0.49, *P* =0.78). Prevalences were also similar in both sexes (χ*^2^_1_*=<0.001, *P* =1.0) and the 3 age classes (χ*^2^_2_*=0.78, P =0.68).

**Table 1.** Prevalence of TBEV RNA and anti-TBEV IgG antibodies in bank voles stratified by trapping site, sex, and age class. Prevalence is presented as percentage with 95% confidence intervals, followed by the number of positive animals and total number tested in brackets.

| Factor | Category | N | PCR+ prevalence, % [95% CI] (n/N) | IgG+ prevalence, % [95% CI] (n/N) |
| --- | --- | --- | --- | --- |
| All animals | - | 258 | 14.3 [10.6-19.1] (37/258) | 6.6 [4.2-10.3] (17/258) |
| Site | Urwitait | 89 | 14.6 [8.7-23.5] (13/89) | 0.0 [0.0-4.1] (0/89) |
|  | Talty | 93 | 16.1 [10.0-24.8] (15/93) | 14.0 [8.4-22.3] (13/93) |
|  | Pilchy | 76 | 11.8 [6.3-21.0] (9/76) | 5.3 [2.1-13.0] (4/76) |
| Sex | Male | 143 | 14.7 [9.9-21.3] (21/143) | 5.6 [2.9-10.7] (8/143) |
|  | Female | 115 | 13.9 [8.7-21.5] (16/115) | 7.8 [4.2-14.2] (9/115) |
| Age class | 1, juvenile | 85 | 11.8 [6.5-20.5] (10/85) | 2.4 [0.7-8.2] (2/85) |
|  | 2, young adult | 118 | 16.1 [10.6-23.7] (19/118) | 7.6 [4.1-13.7] (9/118) |
|  | 3, adult | 55 | 14.5 [7.6-25.7] (8/55) | 10.9 [5.1-21.9] (6/55) |

Serological analysis identified anti-TBEV IgG antibodies in 17 individuals (6.6%; 95% CI: 4.2-10.3). As with PCR positivity, the prevalence of IgG positivity for TBEV did not differ significantly between the sexes (χ*^2^_1_*=<0.001, *P* =0.99) nor between the 3 age classes (χ*^2^_2_*=4.83, P =0.089). Although the association was not statistically significant, IgG prevalence showed an increasing trend from age class 1 to age class 3 (Table 1). There was a highly significant difference in prevalence of IgG positivity between voles from the three sites. No voles were positive for IgG to TBEV in Urwitałt and the highest value was recorded among voles from Tałty. Comparison of molecular and serological results revealed limited concordance between infection and antibody status. Of the 37 RNA-positive individuals, only 9 (24.3%) were also seropositive, whereas 8 individuals were antibody-positive but RNA-negative (Table 1).

### Phylogenetic analysis

Out of 37 TBEV RNA-positive rodent brain samples, all were successfully sequenced for a fragment of the capsid (C) gene, and near-complete genomes were obtained from 10 samples (Table 2). The newly generated sequences were deposited in GenBank under accession numbers PZ361876-PZ361885.

**Table 2.** Pairwise nucleotide and deduced amino acid sequence identities among near-complete TBEV genomes. Values below the diagonal represent nucleotide identity, and values above the diagonal represent deduced amino acid identity. The Neudoerfl strain (GenBank accession U27495) was included as the TBEV-Eu reference.

|  | Neudoerfl<br>_TBEV-Eu | 103_Ta<br>lty_Poland_2018 | 129_Ta<br>lty_Poland_2018 | 130_Ta<br>lty_Poland_2018 | 128_Ta<br>lty_Poland_2018 | 82_Tal<br>ty_Poland_2018 | 162_Ta<br>lty_Poland_2018 | 137_Ta<br>lty_Poland_2018 | 90_Tal<br>ty_Poland_2018 | 153_Ta<br>lty_Poland_2018 | 20_Urw<br>italt_Poland_2018 |
| --- | --- | --- | --- | --- | --- | --- | --- | --- | --- | --- | --- |
| Neudoerfl_TBEV-Eu |  | 99.132 | 99.132 | 99.104 | 99.132 | 99.188 | 99.188 | 99.188 | 99.025 | 99.026 | 98.944 |
| 103_Tal<br>ty_Poland_2018 | 95.241 |  | 100.000 | 99.972 | 100.000 | 99.888 | 99.888 | 99.888 | 99.255 | 99.198 | 99.172 |
| 129_Tal<br>ty_Poland_2018 | 95.232 | 99.991 |  | 99.972 | 100.000 | 99.888 | 99.888 | 99.888 | 99.255 | 99.198 | 99.172 |
| 130_Tal<br>ty_Poland_2018 | 95.213 | 99.972 | 99.981 |  | 99.972 | 99.860 | 99.860 | 99.860 | 99.226 | 99.169 | 99.144 |
| 128_Tal<br>ty_Poland_2018 | 95.232 | 99.991 | 99.981 | 99.963 |  | 99.888 | 99.888 | 99.888 | 99.255 | 99.198 | 99.172 |
| 82_Tal<br>ty_Poland | 95.241 | 99.851 | 99.861 | 99.842 | 99.842 |  | 100.000 | 100.000 | 99.369 | 99.312 | 99.287 |
| d_2018 |  |  |  |  |  |  |  |  |  |  |  |
| 162_Tal<br>ty_Pola<br>nd_201<br>8 | 95.2<br>32 | 99.898 | 99.888 | 99.870 | 99.888 | 99.954 |  | 100.00<br>0 | 99.369 | 99.312 | 99.287 |
| 137_Tal<br>ty_Pola<br>nd_201<br>8 | 95.2<br>32 | 99.898 | 99.888 | 99.870 | 99.888 | 99.954 |  |  | 99.369 | 99.312 | 99.287 |
| 90_Talt<br>y_Polan<br>d_2018 | 95.1<br>19 | 97.859 | 97.850 | 97.840 | 97.850 | 97.878 | 97.869 | 97.869 |  | 99.942 | 99.739 |
| 153_Tal<br>ty_Pola<br>nd_201<br>8 | 95.1<br>25 | 97.824 | 97.814 | 97.795 | 97.814 | 97.833 | 97.824 | 97.824 | 99.981 |  | 99.685 |
| 20_Urw<br>italt_Pol<br>and_20<br>18 | 95.1<br>61 | 97.851 | 97.841 | 97.822 | 97.841 | 97.860 | 97.851 | 97.851 | 99.721 | 99.696 |  |

Phylogenetic analysis based on near-complete genome sequences demonstrated that all isolates clustered within the European subtype (TBEV-Eu), sharing 95.12-95.24% nucleotide identity with the Neudoerfl reference strain (GenBank: U27495) (Figure 3, Table 2), supporting the circulation of TBEV-Eu at the study sites.

**Figure 3.**
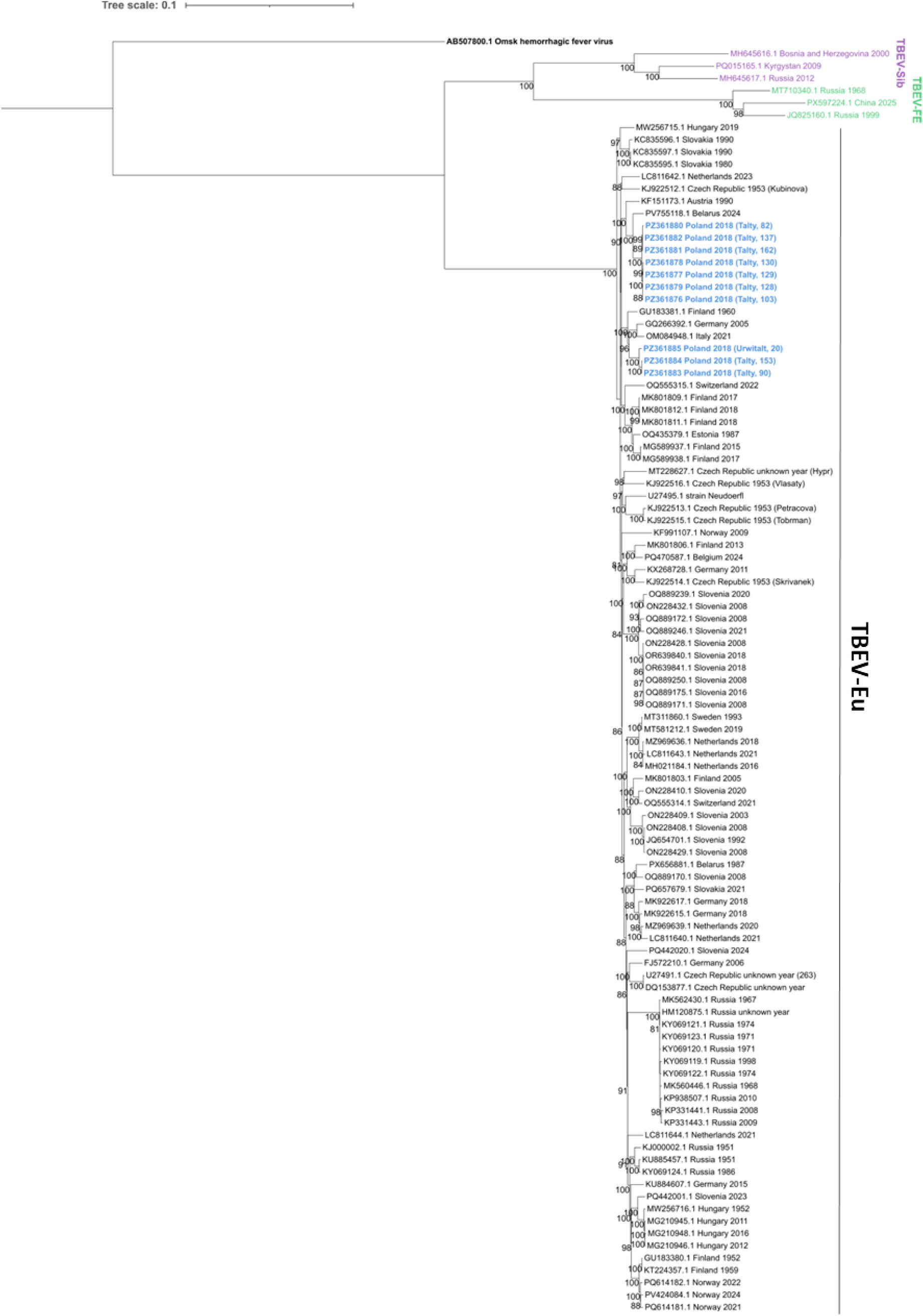
Maximum-likelihood phylogeny of near-complete TBEV genome sequences. Maximum-likelihood tree inferred from full-length TBEV sequences with Omsk hemorrhagic fever virus (GenBank: AB507800) as outgroup. Taxon labels indicate GenBank accession number, country, and collection year. Isolates from this study are in blue and additionally display sampling location and identifier. Bootstrap support values (≥80%) are shown at nodes. Duplicate sequences from identical geographic and temporal origins were removed prior to tree inference. Scale bar, 0.1 substitutions per site.

Despite being collected within a narrow spatial (∼12 km) and temporal (two-week) window, the sequences did not form a single monophyletic group. Instead, two well-supported clades were identified. Nucleotide identity among the obtained sequences ranged from 97.80% to 100% (Table 2).

The first clade comprised seven highly similar sequences from Tałty (samples 82, 103, 128, 129, 130, 137, 162; GenBank accessions: PZ361876-PZ361882). These sequences shared 99.84-100.00% nucleotide identity, with several genomes being identical, and clustered with isolates detected in Belarus (human, 2024) and Austria (*Apodemus flavicollis*, 1990). The second clade included two Tałty isolates (samples 90 and 153; PZ361883-PZ361884) together with the Urwitałt isolate (sample 20; PZ361885). These sequences shared 99.70-99.98% nucleotide identity and grouped with isolates from Germany (*I. ricinus*, 2005), Italy (*Capreolus capreolus*, 2021) and Joutseno strain from Finland (1960).

Amino acid sequence identity among the isolates ranged from 99.14% to 100.00% (Table 2). The seven-isolate Tałty cluster was highly homogeneous, sharing >99.86% amino acid identity, with three isolates exhibiting identical amino acid sequences. The second clade showed slightly greater amino acid variability, with pairwise identities ranging from 99.70% to 99.94%.

### Detection of TBEV RNA and envelope antigen in bank vole embryos

Among TBEV-positive and/or seropositive rodents, five female bank voles were identified as pregnant (Table 3). Three dams were positive for both TBEV RNA and anti-TBEV IgG antibodies, whereas two were seropositive but RNA-negative, consistent with previous infection and the absence of detectable viral RNA in the brain.

**Table 3.** Detection of TBEV RNA and envelope (E) protein in pregnant bank voles and their embryos.

| Dam ID | Dam TBEV IgG | Dam TBEV RNA | Embryos TBEV antigen | Embryos TBEV RNA | Number of embryos | Average embryo length | Average embryo weight [g] |
| --- | --- | --- | --- | --- | --- | --- | --- |

|  |  |  |  |  |  | [cm] |  |
| --- | --- | --- | --- | --- | --- | --- | --- |
| 89 | + | - | + | - | 5 | 0.226 | 0.015 |
| 94 | + | - | - | + | 5 | 2.436 | 1.589 |
| 103 | + | + | + | + | 7 | 1.830 | 1.447 |
| 137 | + | + | + | - | 3 | 0.257 | 0.016 |
| 162 | + | + | + | + | 5 | 0.613 | 0.11 |

To assess possible evidence for vertical transmission, embryos were screened for TBEV RNA using nested RT-PCR. Viral RNA was detected in 3 of 5 litters (60.0%; 95% CI: 23.1-88.2). In addition, embryo-derived homogenates were analyzed for the presence of TBEV envelope (E) protein by Western blot, which was detected in 4 of 5 litters (80.0%; 95% CI: 37.6-96.4) (Figure 4).

**Figure 4.**
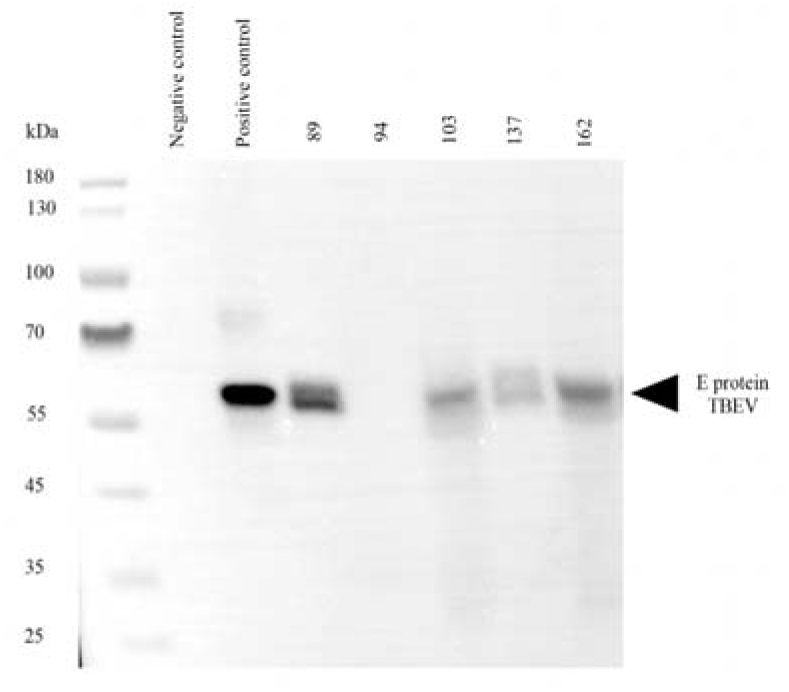
Detection of tick-borne encephalitis virus (TBEV) envelope (E) protein in pooled embryo homogenates from naturally infected bank voles by Western blot. Protein extracts prepared from pooled embryo homogenates from individual litters were separated on an 8% Bolt Bis-Tris SDS–PAGE gel under reducing conditions, transferred to PVDF membranes, and probed with a monoclonal antibody against the TBEV envelope (E) protein. Lane 1, prestained molecular-weight marker; lane 2, negative control (lysate from non-transfected FreeStyle™ 293 cells); lane 3, positive control (lysate from FreeStyle™ 293 cells expressing TBEV E protein); lanes 4–8, pooled embryo homogenates from litters of dams 89, 94, 103, 137, and 162, respectively. TBEV E protein was detected in four of the five litter pools.

Detection patterns varied among litters (Table 3). Two litters from TBEV RNA-positive dams showed concurrent detection of viral RNA and envelope antigen, providing complementary evidence consistent with transplacental exposure or transmission. Litters from seropositive, RNA-negative females exhibited discordant detection patterns, with one litter testing antigen-positive but RNA-negative and another RNA-positive but antigen-negative. Notably, the two RNA-negative litters were the smallest and presumably youngest, suggesting that viral loads in early-stage embryos may fall below detection thresholds or that infection occurs later during gestation. RNA-positive embryos were also detected in one RNA-negative female (No. 94). As maternal RNA testing was performed only on brain tissue, this finding may reflect tissue-specific or temporal differences in viral persistence rather than the absence of infection in the mother. Overall, our findings provide evidence for the presence of TBEV RNA and/or viral antigen in embryos from naturally infected wild rodents.

## Discussion

According to the latest ECDC Annual Epidemiological Report (2022 data), the incidence of tick-borne encephalitis and its geographic range are expanding across Europe, with a 14% increase in 2022 compared with 2021, continuing the upward trend observed since 2012 [45]. Similar patterns are evident in Poland, where case numbers are growing each year and infections are increasingly reported outside historically endemic regions in the northeast [46]. This expansion likely reflects climatic, ecological, and behavioral changes, highlighting the need for a deeper understanding of TBEV maintenance mechanisms in natural systems and the distribution of circulating TBEV lineages [45, 47, 48].

Despite Poland’s proximity to regions where TBEV subtypes co-circulate [49], all sequences obtained in this study belonged to the European subtype (TBEV-Eu), supporting previous evidence that TBEV-Eu remains the dominant subtype in Poland [24, 25, 50, 51]. The absence of TBEV-Sib and TBEV-FE may reflect ecological barriers or vector distribution limits. However, the known range of *I. persulcatus* extends increasingly close to Polish borders, highlighting the need for continued genetic monitoring, as introduction of more virulent subtypes could alter disease patterns[22, 52–54].

TBEV sequences from a small geographic area segregated into two distinct clades, suggesting complex local transmission dynamics consistent with previous observations of phylogenetically distinct lineages co-circulating within geographically restricted foci [33]. Whether these lineages occupy distinct ecological micro-foci or represent transient co-circulation remains unclear and warrants further longitudinal investigation.

The discrepancy between TBEV RNA detection and IgG seroprevalence likely reflects differences between experimental and natural infection trajectories, as well as the tissues sampled. Natural infections are more heterogeneous, with wild rodents sampled at unknown infection stages. This is consistent with previous findings showing RNA-positive but antibody-negative bank voles in TBEV-Eu endemic areas [55]. Moreover, TBEV RNA has been detected in wild rodents outside the tick-feeding season, indicating that persistent infection may contribute to virus overwintering [55], and experimental evidence confirms viral RNA may persist in brain tissue for extended periods post-infection [15]. Morozova et al. (2020) further showed that viral RNA may persist in wild rodents despite the presence of specific antibodies, supporting long-term TBEV persistence in reservoir hosts [56]. As TBEV can replicate locally within the epidermis without systemic dissemination [57], our brain-tissue-based prevalence estimates may be conservative [58]. These findings indicate that seroprevalence alone may substantially underestimate true infection rates in natural populations.

While vertical transmission has been demonstrated experimentally for other subtypes and closely related viruses [12, 59], and documented in natural reservoir populations for TBEV-FE [13], evidence from wild systems for TBEV-Eu remains absent. Transmission was not uniform across litters, with discordant RNA and antigen detection patterns suggesting heterogeneity related to infection timing, maternal viral load, or embryo developmental stage. The smallest litters were RNA-negative, possibly reflecting detection thresholds or timing of infection during gestation.

The ecological implications of vertical transmission are potentially substantial. Unlike incidental hosts with short viremia, reservoir hosts may support persistence through multiple complementary transmission routes. Vertical transmission may contribute to virus circulation during periods of low tick activity, stabilizing transmission cycles across seasons. Combined with co-feeding transmission and persistent infections, this pathway may facilitate long-term TBEV persistence in localized natural foci.

Several limitations must be acknowledged. We could not determine whether infected embryos survive to birth or contribute to onward transmission, and viral sequencing from embryo samples was unsuccessful, likely due to low viral loads.

In summary, this study provides the first near-complete genome sequences of TBEV from rodents in Poland, confirms circulation of the European subtype, and provides the first evidence consistent with vertical transmission of TBEV-Eu in a natural reservoir host. Together with local co-circulation of distinct viral lineages and limited concordance between RNA detection and serology, these findings highlight the heterogeneous and multifaceted nature of TBEV infection dynamics under natural conditions. Future studies should focus on longitudinal monitoring of infection markers and offspring from infected dams, extending beyond the snapshot provided by the present study. They should also experimentally validate vertical transmission efficiency and quantify its relative contribution to virus persistence compared with vector-mediated routes. Expanded ecological and molecular surveillance will be essential to understand virus maintenance in natural systems and to develop effective strategies for mitigating the impact of TBEV in both endemic and emerging areas.

## ETHICS APPROVAL

This study was carried out according to the recommendations in the Guidelines for the Care and Use of Laboratory Animals of the Polish National Ethics Committee for Animal Experimentation. The study was performed according to the ARRIVE guidelines 2.0.

## AVAILABILITY OF DATA AND MATERIALS

The newly generated TBEV genome sequences have been deposited in the NCBI GenBank database under accession numbers PZ361876–PZ361885. Raw sequencing data are available through the NCBI BioProject database under accession number PRJNA953225, with associated BioSample accession numbers SAMN35084469–SAMN35084478. The individual-level metadata underlying the prevalence, seroprevalence, and ecological analyses have been deposited in the Zenodo repository and are publicly available at https://doi.org/10.5281/zenodo.21532124.

## CONSENT FOR PUBLICATION

Not Applicable

## AUTHORS’ CONTRIBUTIONS

The study was conceived and designed by MG, MK, OV, TSir. Supervision of the long-term monitoring of bank vole populations in the region was by JMB, AB and MG. Samples were collected and dissected in the field by JMB, AB, MG, BB, KT and DD. The laboratory work was conducted by MK, TS, KBar and RK. Immunological analyses - SM, LR, KBas and MKR. Molecular analyses - MK, TS, RK and KBar. Data handling - MG. Statistical analyses were carried out by MG, MK and JMB. The manuscript was written by MK, MG and JMB in consultation with all co-authors. MG & MK revised the manuscript. Project administration - MG. Funding acquisition - MG, JMB, AB and RK. All authors accepted the final manuscript version.

## ACKNOWLEDGEMENTS

MG thanks Mira Utriainen and Ewa Zieliniewicz for their assistance with the laboratory work.

## REFERENCES

1. Lindquist L, Vapalahti O. Tick-borne encephalitis. The Lancet 2008;371:1861–1871. 10.1016/S0140-6736(08)60800-4

2. Bogovic P, Lotric-Furlan S, Strle F. What tick-borne encephalitis may look like: clinical signs and symptoms. Travel Med Infect Dis 2010;8:246–250. 10.1016/j.tmaid.2010.05.011

3. Ecker M, Allison S, Meixner T et al. Sequence analysis and genetic classification of tick-borne encephalitis viruses from Europe and Asia J Gen Virol 1999;80:179–185. 10.1099/0022-1317-80-1-179

4. Gritsun TS, Lashkevich VA, Gould EA et al. Characterization of a Siberian Virus Isolated from a Patient with Progressive Chronic Tick-Borne Encephalitis. J Virol 2003;77:25–36. 10.1128/jvi.77.1.25-36.2003

5. Ruzek D, Avšič Županc T, Borde J et al. Tick-borne encephalitis in Europe and Russia: Review of pathogenesis, clinical features, therapy, and vaccines. Antiviral Res 2019;164:23–51. 10.1016/j.antiviral.2019.01.014

6. Kožuch O, Grešíková M, Nosek J et al. The role of small rodents and hedgehogs in a natural focus of tick-borne encephalitis. Bull World Health Organ 1967;36:61–66.

7. Achazi K, Růžek D, Donoso-Mantke O et al. Rodents as sentinels for the prevalence of tick-borne encephalitis virus. Vector Borne Zoonotic Dis 2011;11:641–647. 10.1089/vbz.2010.0236

8. Stefanoff P, Pfeffer M, Hellenbrand W et al. Virus detection in questing ticks is not a sensitive indicator for risk assessment of tick-borne encephalitis in humans. Zoonoses Public Health 2013;60:215–226. 10.1111/j.1863-2378.2012.01517.x

9. Grzybek M, Cybulska A, Tołkacz K et al. Seroprevalence of TBEV in bank voles from Poland—a long-term approach. Emerg Microbes Infect 2018;7:1–8. 10.1038/s41426-018-0149-3

10. Zöldi V, Papp T, Reiczigel J et al. Bank voles show high seropositivity rates in a natural TBEV focus in Hungary. Infect Dis 2015;47:178–181. 10.3109/00365548.2014.975743

11. Pintér R, Madai M, Horváth G et al. Molecular detection and phylogenetic analysis of tick-borne encephalitis virus in rodents captured in the transdanubian region of Hungary. Vector Borne Zoonotic Dis 2014;14:621–624. 10.1089/vbz.2013.1479

12. Gerlinskaya LA, Bakhvalova VN, Morozova OV. Sexual transmission of tick-borne encephalitis virus in laboratory mice. Bull Exp Biol Med 1997;123:283–284. 10.1007/BF02445427

13. Bakhvalova VN, Potapova OF, Panov VV et al. Vertical transmission of tick-borne encephalitis virus between generations of adapted reservoir small rodents. Virus Res 2009;140:172–178. 10.1016/j.virusres.2008.12.001

14. Tonteri E, Kurkela, S., Timonen, S, et al. Surveillance of endemic foci of tick-borne encephalitis in Finland 1995–2013: evidence of emergence of new foci. Euro Surveill 2015;20:30020. 10.2807/1560-7917.ES.2015.20.37.30020

15. Tonteri E, Kipar A, Voutilainen L et al. The three subtypes of tick-borne encephalitis virus induce encephalitis in a natural host, the bank vole (Myodes glareolus). PLoS One 2013;8:e81214. 10.1371/journal.pone.0081214

16. Topp AK, Springer A, Dobler G et al. New and confirmed foci of tick-borne encephalitis virus (TBE) in northern Germany determined by TBEV detection in ticks. Pathogens 2022;11:126. 10.3390/pathogens11020126

17. Dekker M, Laverman GD, de Vries A et al. Emergence of tick-borne encephalitis (TBE) in the Netherlands. Ticks Tick Borne Dis 2019;10:176–179. 10.1016/j.ttbdis.2018.10.008

18. Holding M, Dowall S, Hewson R. Detection of tick-borne encephalitis virus in the UK. Lancet 2020;395:411. 10.1016/S0140-6736(20)30040-4

19. Fomsgaard A, Christiansen CB, Bødker R. First identification of tick-borne encephalitis in Denmark outside of Bornholm, August 2009. Euro Surveill 2009;14:19325. 10.2807/ese.14.36.19325-en

20. Andersen NS, Larsen SL, Olesen CR et al. Continued expansion of tick-borne pathogens: Tick-borne encephalitis virus complex and Anaplasma phagocytophilum in Denmark. Ticks Tick Borne Dis 2019;10:115–123. 10.1016/j.ttbdis.2018.09.007

21. Jääskeläinen AE, Tikkakoski T, Uzcátegui NY et al. Siberian subtype tickborne encephalitis virus, Finland. Emerg Infect Dis 2006;12:1568–1571. 10.3201/eid1210.060320

22. Jaenson TGT, Värv K, Fröjdman I et al. First evidence of established populations of the taiga tick Ixodes persulcatus (Acari: Ixodidae) in Sweden. Parasit Vectors 2016;9:377. 10.1186/s13071-016-1658-3

23. Philippe C, De Sterck C, Parys A et al. First detection of tick-borne encephalitis virus in Ixodes ricinus ticks in Belgium, May 2024. Parasit Vectors 2025 18:1 2025;18:197-. 10.1186/S13071-025-06829-5

24. Katargina O, Russakova S, Geller J et al. Detection and characterization of tick-borne encephalitis virus in Baltic countries and eastern Poland. PLoS One 2013;8:e61374. 10.1371/journal.pone.0061374

25. Wójcik-Fatla A, Cisak E, Zając V et al. Prevalence of tick-borne encephalitis virus in Ixodes ricinus and Dermacentor reticulatus ticks collected from the Lublin region (eastern Poland). Ticks Tick Borne Dis 2011;2:16–19. 10.1016/j.ttbdis.2010.10.001

26. Kloch A, Babik W, Bajer A et al. Effects of an MHC-DRB genotype and allele number on the load of gut parasites in the bank vole Myodes glareolus. Mol Ecol 2010;19:255–265. 10.1111/j.1365-294X.2009.04476.x

27. Behnke JM, Barnard CJ, Bajer A et al. Variation in the helminth community structure in bank voles (Clethrionomys glareolus) from three comparable localities in the Mazury Lake District region of Poland. Parasitology 2001;123:401–414. DOI:10.1017/S0031182001008605

28. Behnke JM, Bajer A, Harris PD et al. Temporal and between-site variation in helminth communities of bank voles (Myodes glareolus) from N.E. Poland. 1. Regional fauna and component community levels. Parasitology 2008;135:985–997. DOI: 10.1017/S0031182008004393

29. Behnke JM, Bajer A, Harris PD et al. Temporal and between-site variation in helminth communities of bank voles (Myodes glareolus) from N.E. Poland. 2. The infracommunity level. Parasitology 2008;135:999–1018. DOI: 10.1017/S0031182008004484

30. Morris P. A review of mammalian age determination methods. Mamm Rev 1972;2:69–104. 10.1111/j.1365-2907.1972.tb00160.x

31. Ramelow C, Süss J, Berndt D et al. Detection of tick-borne encephalitis virus RNA in ticks (Ixodes ricinus) by the polymerase chain reaction. J Virol Methods 1993;45:115–119. 10.1016/0166-0934(93)90145-H

32. Kallio-Kokko H, Laakkonen J, Rizzoli A et al. Hantavirus and arenavirus antibody prevalence in rodents and humans in Trentino, northern Italy. Epidemiol Infect 2006;134:830–836. 10.1017/S0950268805005431

33. Smura T, Tonteri E, Jääskeläinen A et al. Recent establishment of tick-borne encephalitis foci with distinct viral lineages in the Helsinki area, Finland. Emerg Microbes Infect 2019;8:675–683. 10.1080/22221751.2019.1612279

34. Chen S, Zhou Y, Chen Y et al. fastp: an ultra-fast all-in-one FASTQ preprocessor. Bioinformatics 2018;34:i884–i890. 10.1093/bioinformatics/bty560

35. Li H. Aligning sequence reads, clone sequences and assembly contigs with BWA-MEM. arXiv preprint 2013. *arXiv:1303.*3997

36. Truong Nguyen PT, Plyusnin I, Sironen T et al. HAVoC, a bioinformatic pipeline for reference-based consensus assembly and lineage assignment for SARS-CoV-2 sequences. BMC Bioinformatics 2021;22:373. 10.1186/S12859-021-04294-2

37. Katoh K, Standley DM. MAFFT Multiple Sequence Alignment Software Version 7: Improvements in Performance and Usability. Mol Biol Evol 2013;30:772–780. 10.1093/molbev/mst010

38. Minh BQ, Schmidt HA, Chernomor O et al. IQ-TREE 2: new models and efficient methods for phylogenetic inference in the genomic era. Mol Biol Evol 2020;37:1530–34. 10.1093/molbev/msaa015

39. Kalyaanamoorthy S, Minh BQ, Wong TKF et al. ModelFinder: fast model selection for accurate phylogenetic estimates. Nat Methods 2017;14:587–589. 10.1038/nmeth.4285

40. Hoang DT, Chernomor O, von Haeseler A et al. UFBoot2: Improving the Ultrafast Bootstrap Approximation. Mol Biol Evol 2018;35:518–522. 10.1093/MOLBEV/MSX281

41. Bajer A, Behnke JM, Pawełczyk A et al. Medium-term temporal stability of the helminth component community structure in bank voles (Clethrionomys glareolus) from the Mazury Lake District region of Poland. Parasitology 2005;130:213–228. 10.1017/S0031182004006389

42. Grzybek M, Bajer A, Bednarska MM et al. Long-term spatiotemporal stability and dynamic changes in helminth infracommunities of bank voles (Myodes glareolus) in NE Poland. Parasitology 2015;142:1722–1743. 10.1017/S0031182015001225

43. Rohlf FJ, Sokal RR. Statistical tables. Macmillan 1995.

44. Percie du Sert N, Hurst V, Ahluwalia A et al. The ARRIVE guidelines 2.0: updated guidelines for reporting animal research. J Cereb Blood Flow Metab 2020;40:1769–77. 10.1177/0271678X20943823

45. European Centre for Disease Prevention and Control. Tick-borne encephalitis: annual epidemiological report for 2022. Stockholm: ECDC, 2024. https://www.ecdc.europa.eu/en/publications-data/tick-borne-encephalitis-annual-epidemiological-report-2022

46. National Institute of Public Health NIH – National Research Institute. Infectious diseases and poisonings in Poland – 2025. https://wwwold.pzh.gov.pl/oldpage/epimeld/2025/index_mp.html (3 May 2026, date last accessed).

47. Stefanoff P, Rosińska M, Samuels S et al. A national case-control study identifies human socio-economic status and activities as risk factors for tick-borne encephalitis in Poland. PLoS One 2012;7:e45511. 10.1371/journal.pone.0045511

48. Maqbool M, Rana A, Khan H, et al. Impact of climate change on tick-borne viral diseases. In: Rizwan HM, Sajid MS (eds), Ticks in a Changing Climate. Cham: Springer, 2026, 85–107. 10.1007/978-3-032-12259-9_5

49. Lundkvist Å, Vene S, Golovljova I et al. Characterization of tick-borne encephalitis virus from Latvia: evidence for co-circulation of three distinct subtypes. J Med Virol 2001;65:730–735. 10.1002/jmv.2097

50. Król N, Chitimia-Dobler L, Dobler G et al. Identification of new microfoci and genetic characterization of tick-borne encephalitis virus isolates from eastern Germany and western Poland. Viruses 2024;16:637. 10.3390/V16040637

51. Biernat B, Karbowiak G, Werszko J et al. Prevalence of tick-borne encephalitis virus (TBEV) RNA in *Dermacentor reticulatus* ticks from natural and urban environment, Poland. Exp Appl Acarol 2014;64:543–551. 10.1007/s10493-014-9836-5

52. Bugmyrin SV, Romanova LY, Belova OA et al. Pathogens in Ixodes persulcatus and Ixodes ricinus ticks (Acari, Ixodidae) in Karelia (Russia). Ticks Tick Borne Dis 2022;13:102045. 10.1016/j.ttbdis.2022.102045

53. Golovljova I, Vene S, Sjölander KB et al. Characterization of tick-borne encephalitis virus from Estonia. J Med Virol 2004;74:580–588. 10.1002/jmv.20224

54. Kuivanen S et al. Fatal Tick-Borne Encephalitis Virus Infections Caused by Siberian and European Subtypes, Finland, 2015. Emerg Infect Dis 2018;24:946. 10.3201/EID2405.171986

55. Kuivanen S, Smura T, Rantanen K et al. Tick-borne encephalitis virus in wild rodents in winter, Finland, 2008-2009. *Emerg Infect Dis* 2011;17:72–75. 10.3201/eid1701.100051

56. Morozova O V., Panov V V., Bakhvalova VN. Innate and adaptive immunity in wild rodents spontaneously and experimentally infected with the tick-borne encephalitis virus nfect Genet Evol 2020;80:104187. 10.1016/J.MEEGID.2020.104187

57. Labuda M, Kozuch O, Zuffová E et al. Tick-borne encephalitis virus transmission between ticks cofeeding on specific immune natural rodent hosts. Virology 1997;235:138–143. 10.1006/viro.1997.8622

58. Pascoe EL, de Vries A, Esser HJ et al. Detection of tick-borne encephalitis virus in ear tissue and dried blood spots from naturally infected wild rodents. Parasit Vectors 2023;16:103. 10.1186/s13071-023-05717-0

59. Miao Y, Zheng Y, Wang T et al. Breast milk transmission and involvement of mammary glands in tick-borne flavivirus infected mice. J Virol 2024;98. 10.1128/jvi.01709-23

